# Deep mutational scanning comprehensively maps how Zika envelope protein mutations affect viral growth and antibody escape

**DOI:** 10.1101/725556

**Authors:** Marion Sourisseau, Daniel J.P. Lawrence, Megan C. Schwarz, Carina H. Storrs, Ethan C. Veit, Jesse D. Bloom, Matthew J. Evans

**Author notes:** These authors contributed equally to this work (Marion Sourisseau, Daniel J.P. Lawrence). Corresponding authors: Jesse D. Bloom, Ph.D., Matthew J. Evans, Ph.D.

## Abstract

Functional constraints on viral proteins are often assessed by examining sequence conservation among natural strains, but this approach is relatively ineffective for Zika virus because all known sequences are highly similar. Here we take an alternative approach to map functional constraints on Zika virus’s envelope (E) protein by using deep mutational scanning to measure how all amino-acid mutations to the protein affect viral growth in cell culture. The resulting sequence-function map is consistent with existing knowledge about E protein structure and function, but also provides insight into mutation-level constraints in many regions of the protein that have not been well characterized in prior functional work. In addition, we extend our approach to completely map how mutations affect viral neutralization by two monoclonal antibodies, thereby precisely defining their functional epitopes. Overall, our study provides a valuable resource for understanding the effects of mutations to this important viral protein, and also offers a roadmap for future work to map functional and antigenic selection to Zika virus at high resolution.

**Importance:** Zika virus has recently been shown to be associated with severe birth defects. The virus’s E protein mediates its ability to infect cells, and is also the primary target of the antibodies that are elicited by natural infection and vaccines that are being developed against the virus. Therefore, determining the effects of mutations to this protein is important for understanding its function, its susceptibility to vaccine-mediated immunity, and its potential for future evolution. We completely mapped how amino-acid mutations to E protein affected the virus’s ability to grow in cells in the lab and escape from several antibodies. The resulting maps relate changes in the E protein’s sequence to changes in viral function, and therefore provide a valuable complement to existing maps of the physical structure of the protein.

## INTRODUCTION

Zika virus (ZIKV) became the subject of intense interest after recent human outbreaks in the Yap Islands, French Polynesia, and Brazil^1–3^ were associated with severe birth defects and neurological disease^4,5^. ZIKV is a member of the genus *Flavivirus*, and is closely related to the dengue, West Nile, yellow fever, and Japanese encephalitis viruses^6^. Like all viruses in this genus, ZIKV has an approximately 11.8-kilobase capped positive-sense, single-stranded RNA genome. This RNA encodes a single open reading frame that is translated into an approximately 3,432 amino-acid polyprotein that is processed by host and viral proteases into three structural and seven non-structural proteins. The mature infectious virion is comprised of a single copy of the RNA genome, surrounded by a capsid (C) protein shell and a lipid envelope bearing 180 copies each of the membrane (M) and envelope (E) proteins^7^.

Immature ZIKV particles bearing 60 heterotrimeric premembrane (prM) and E protein spikes assemble at the endoplasmic reticulum and transit through the trans-Golgi network, where E proteins undergo rearrangement into anti-parallel dimers arranged in a herringbone pattern characteristic of flaviviruses^7^. This rearrangement exposes the prM cleavage site that is cleaved by a furin-like protease. Upon release of the virion into the more neutral pH of the extracellular environment, pr dissociates from the M and E protein, resulting in the mature virion. The E protein in the mature virion is folded into three domains^8,9^. Domain I (DI) is comprised of a central β-barrel and a highly flexible loop that bears the sole E protein N-linked glycosylation site, N154. This glycan is highly conserved between most flaviviruses, but is absent from some ZIKV isolates, particularly those of the African lineage^10,11^. Domain II (DII) forms an elongated ‘finger-like’ shape that is the major interaction surface between monomers of an E dimer. The fusion loop is located at the region of DII that is distal to domains I and III. The major feature of domain III (DIII) is an immunoglobulin-like region that is thought to contribute to host cell binding^12,13^. However, an E protein binding receptor has not yet been identified and thus the viral determinants for such interactions are not defined.

In order to maintain viral fitness, the E protein must retain the capacity to undergo the complex series of protein-protein interactions and conformational changes to form mature viral particles. Once these viral particles are released from a cell, the E protein needs to functionally interact with both insect and mammalian target cells in order to mediate viral binding and fusion. At the same time, the E protein is the primary target of neutralizing antibodies. As such, this protein is subjected to an array of evolutionary pressures. Any viral antigenic variation is expected to be due largely to mutations in the E protein. For many viral proteins, it is possible to identify functionally constrained motifs and sites with capacity for evolutionary change by examining conservation in alignments of natural viral variants. However, because ZIKV has only been carefully studied very recently, nearly all available sequences are from recent viruses. Furthermore, simply comparing viral sequences shows little differences across both older African and more recent Asian lineages, with even the most divergent protein sequences >95% identical^14^. Inspection of natural sequences is therefore not an effective method to identify constraint and selection on E protein.

An alternative way to assess the mutational constraints on a protein is to use deep mutational scanning to measure the effects of all amino-acid mutations to a protein on viral growth in cell culture and antigenic recognition by antibodies. Deep mutational scanning is a relatively new approach that involves creating large libraries of mutants of a gene, imposing a functional selection, and using deep sequencing to estimate the effect of every mutation from its change in frequency after selection^15^. This approach has been used to assess the effects of all viral mutations on the function and antigenicity of key proteins from influenza virus^16–18^ and HIV^19,20^. Two studies have applied deep mutational scanning to ZIKV^21,22^, but these studies did not systematically map the mutational tolerance or antigenicity of all amino-acid mutations. Instead, they used libraries comprised of a subset of E protein variants to identify mutations that differentially affect viral growth in mammalian and insect cells.

Here we apply deep mutational scanning to measure the effects of all amino-acid mutations to the ZIKV E protein on viral growth in cell culture. The resulting sequence-function map identifies which sites in E protein are amenable to mutational change, and which ones are under functional constraint. In addition, we completely map mutations that alter the antigenicity of E protein with respect to antibodies targeting two distinct regions. Overall, our results provide a map of the fine-grained constraints on E protein and demonstrate a powerful platform for further functional and antigenic investigations of this protein.

## RESULTS

### Deep mutational scanning to measure the effects of all amino-acid mutations to ZIKV E protein on viral growth in cell culture

To measure the effects of all amino-acid mutations to the ZIKV E protein, we followed the experimental scheme illustrated in Fig 1A. First, we used a previously developed PCR-based technique^23,24^ to create a library of all codon mutants of the full-length E protein-coding gene. We created this library in our previously reported infectious clone of the ZIKV prototype strain MR766^25^, choosing this clone because the plasmid is stable in bacteria (which facilitated the construction of the mutant library) and because the virus grows to high titer (which helped avoid bottlenecks during generation of the mutant virus library). We created three independent E protein mutant libraries, which were handled fully independently in all subsequent steps to provide true biological replicates (Fig 1B). Each library replicate was comprised of >10^6^ unique plasmid transformants. There are 504 codons in the MR766 E gene, thus with 19 alternative amino acids there are 9,576 possible amino-acid mutations and 31,752 possible mutant codon sequences if all nucleotide positions in each codon are randomized. Deep sequencing the plasmid library showed that virtually all of these mutations were observed multiple times in each replicate library (Fig 1C). Our mutagenesis method is expected to introduce a Poisson number of codon mutations per variant^23,24^, and Sanger sequencing of 40 plasmids verified that the number of mutations per variant followed a roughly Poisson distribution with a mean of 1.7 (Fig 1D), meaning that our libraries measure the average effect of each amino-acid mutation alone and in several closely related backgrounds.

**FIG 1:**
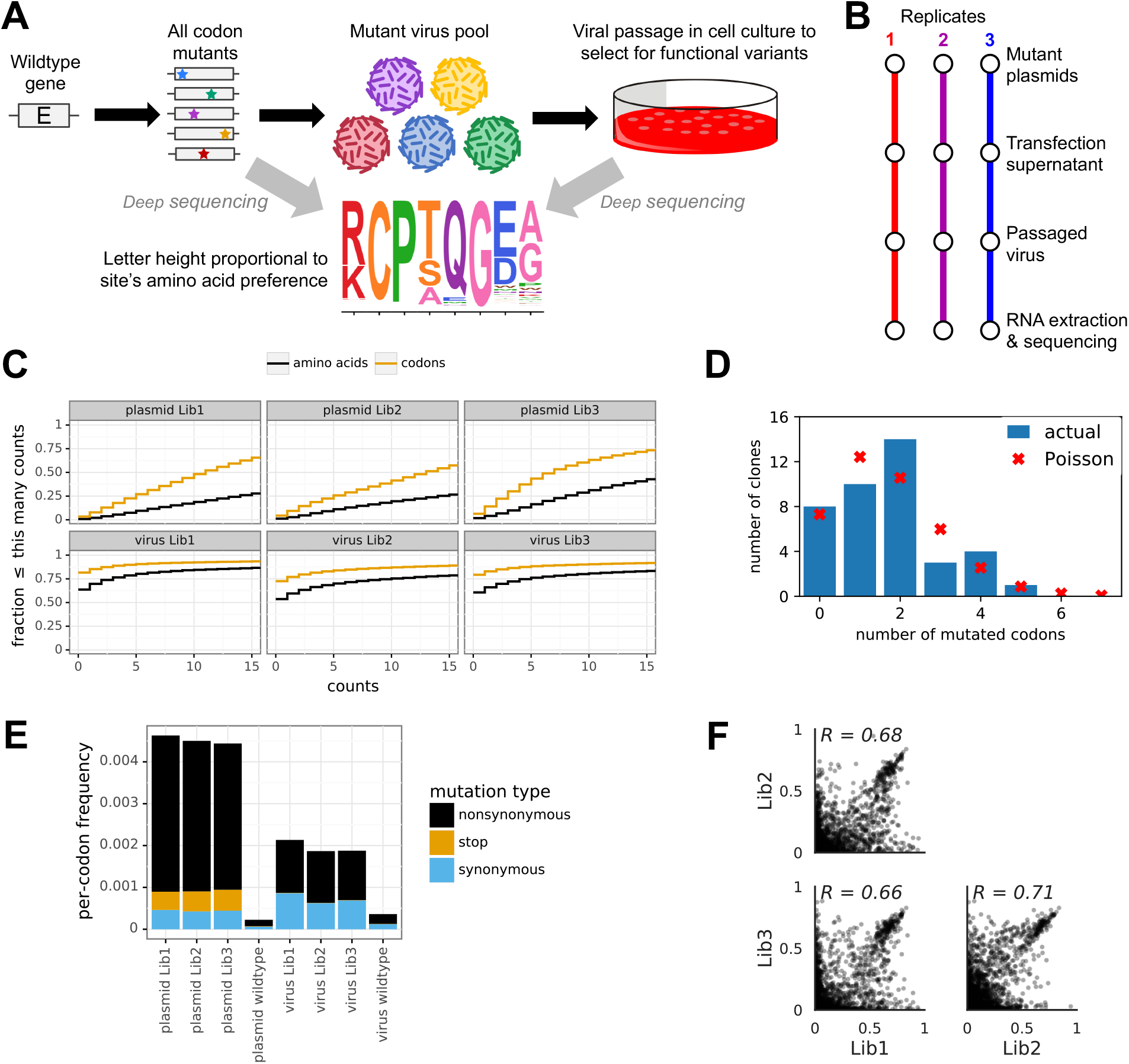
Deep mutational scanning of ZIKV E protein. **(A)** Overview of experimental approach. We generated plasmid libraries with all codon mutations to the E protein gene, and then used these plasmids to generate libraries of mutant viruses that we passaged in cell culture at low MOI. By using deep sequencing to measure the frequency of each mutation in the initial plasmid library versus the passaged mutant viruses, we determined the amino acid preference of each site. **(B)** The deep mutational scanning was performed in full biological triplicate, with each replicate beginning with independent generation of a mutant library. **(C)** Fraction of all aminoacid or codon mutations observed at least the indicated number of times in the original plasmid libraries and the viruses that have undergone functional selection. **(D)** Sanger sequencing of 40 clones in the plasmid library showed that the number of codon mutations per clone was roughly Poisson distributed. **(E)** Frequency of each type of codon mutation averaged across the entire gene in the original plasmid libraries, the passaged virus libraries, and wild-type plasmid and virus controls. **(F)** The correlations between the amino-acid preferences measured in each biological replicate. Each point represents a preference for a specific amino acid at a specific site, and the numbers show the Pearson correlation coefficient.

To generate mutant viruses from the mutant plasmids (Fig 1A), we transfected 293T cells with the plasmid DNA libraries to produce pools of viruses with genomes encoding all the E protein mutants. Cells transfected with the mutant plasmid libraries produced titers of 9×10^4^ to 5×10^5^ infectious units per mL after 48 hours, which was 240 to 1280-fold less than the titers obtained with the wild-type ZIKV genome. To select for the functional variants in our virus libraries, we infected Vero cells with the transfection supernatants at a multiplicity of infection (MOI) of 0.01, using 10^8^ cells to maintain diversity of 10^6^ independent E protein mutants. A low MOI was required to link the mutant viral genome with its respective virion, as virus produced from transfected 293T cells may have a genotype-phenotype mismatch (i.e., the E protein variants that compose the virion are likely to be mixtures and may not be the same as the one encoded on the packaged genome). Low-MOI infection also imposed a selective barrier: viruses underwent multi-cycle replication in proportion to the functionality of their E protein. At 24 hours post-infection, we washed the infected cells to remove input virus, and then let the infection continue for two additional days. At this point, viral variants with high fitness had undergone 3-4 rounds of additional infection and thus would be enriched in the population.

We then used deep sequencing to quantify the frequency of each mutation in the mutant viruses relative to the initial plasmid mutant libraries. To sequence the passaged viruses, we extracted and reverse-transcribed RNA from the infected Vero cells. In order to ensure high sequencing accuracy, we used a previously described barcoded-subamplicon sequencing approach^16,17^, which uses random nucleotide barcodes attached to template molecules to correct for sequencing errors. As shown in Fig 1E, the initial plasmid libraries had a high rate of mutations, which included nonsynonymous, stop-codon, and synonymous mutations. After viral passaging, the number of mutations decreases sharply. In particular, stop codons (which are expected to be uniformly deleterious) were purged, and nonsynonymous mutations (many of which will be deleterious) decreased in frequency. Overall, slightly less than half of all aminoacid mutations were still observed in the mutant viruses after passaging (Fig 1C). Fig 1E also shows that the apparent rate of mutations in wild-type plasmid or virus generated from wild-type plasmid was low, indicating that sequencing, reverse-transcription, and viral replication errors made only modest contributions to the mutations observed in the sequencing.

We used the sequencing data to measure the “preference” of each site in E protein for each amino acid. These preferences represent enrichments of each amino-acid at each site after selection for viral growth, normalized to the abundance of the wild type codon at each site. Fig 1F shows the correlation among all 10,080 amino-acid preference values (there are 504 E protein codons multiplied by 20 unique amino acids for 10,080 preference values) between each biological replicate of the experiment. Although there is some noise, the preferences were strongly correlated among replicates. For the remainder of this paper, we use the average of the three replicates. The across-replicate average of the amino-acid preferences for all sites in E are shown in Fig 2A. Visual inspection of this figure reveals that the wild-type amino acid was usually, but not always, the most preferred amino acid at a site. It is also apparent that mutational tolerance varies widely across the E protein, with some sites strongly preferring a specific amino acid (e.g., site 10) while other sites tolerated many amino acids (e.g., site 36). The amino-acid preference data in Fig 2A can also be mathematically transformed into a measurement of the effect of each mutant amino acid relative to the wild-type identity at that site in MR766. These mutational effects are shown in Fig 2B — as is apparent from inspection of this figure, mutations were usually deleterious (below the black center line), but this was much truer for some sites than others. For instance, most mutations were strongly deleterious at the mutationally intolerant site 10, whereas many mutations only had small effects at the mutationally tolerant site 36. Overall, these maps provide a wealth of information about the sequence-function relationships in E protein.

**FIG 2:**
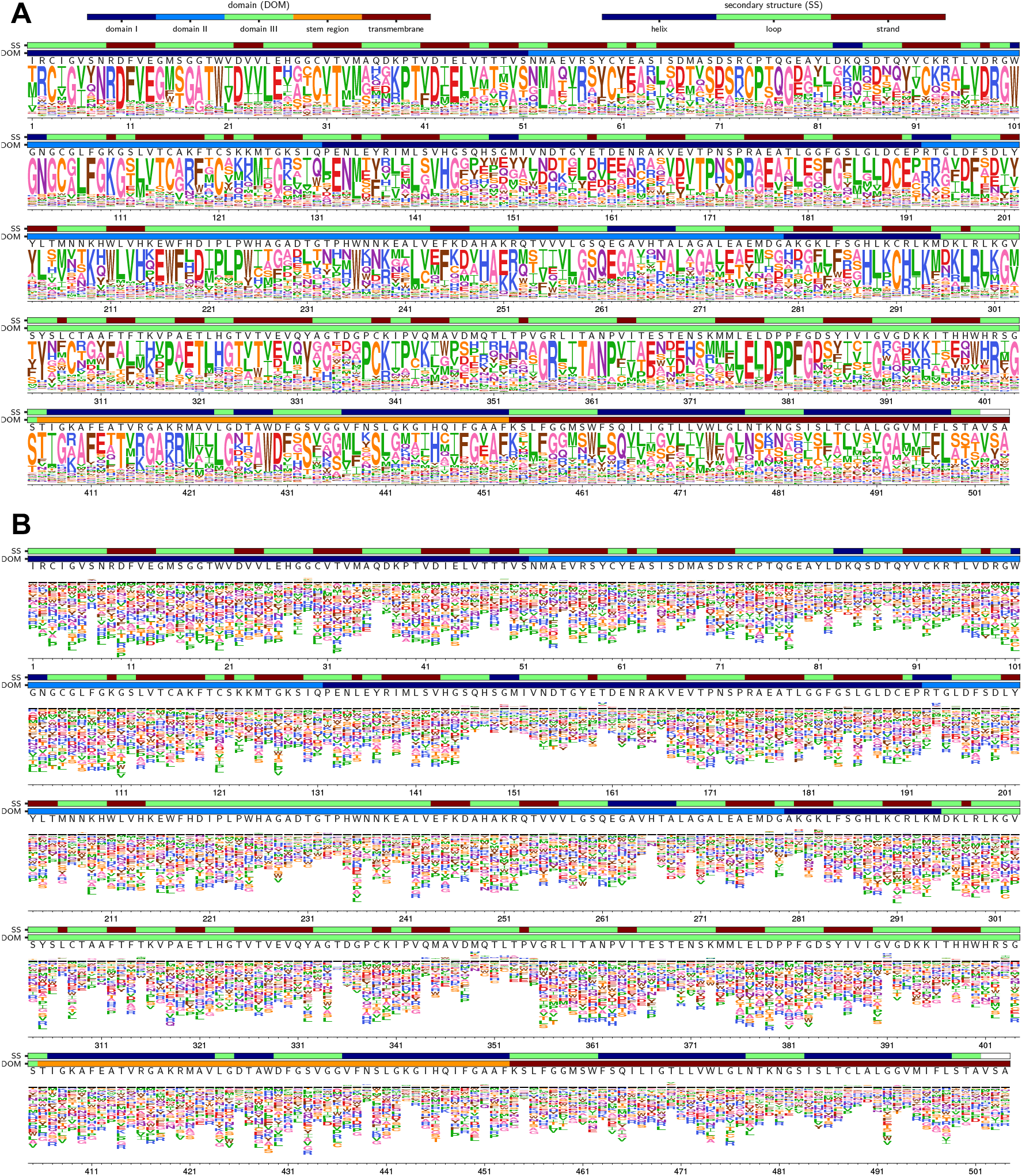
Effects of all amino-acid mutations at each site in E protein as measured in the deep mutational scanning. **(A)** The preference of each site for each amino acid. The height of each letter is proportional to the enrichment of that amino acid after selection relative to its frequency before selection, so taller letters indicate more strongly preferred amino acids. The letters immediately above each logo stack indicate the wild-type amino acid in the parental MR766 E protein, the first color bar (DOM) indicates the domain of E protein as defined in Dai et al.^26^, and the second color bar (SS) indicates the secondary structure of that region as defined using DSSP^48^ on PDB 5ire^7^. **(B)** An alternative representation of the same data shown in panel A. In this representation, the height of each letter represents the estimated effect of that mutation, where letters above the black line indicate favorable mutations (there are relatively few of these) and letters below the black line indicate unfavorable mutations. The effect of a mutant amino acid is equal to the logarithm of the ratio of its preference divided by the preference for the wildtype amino acid at that site.

### Comparing mutational effects measured in the lab to amino-acid frequencies in nature

We next examined how the amino-acid preferences measured in our screen compared to frequencies of amino acids at each site in an alignment of naturally occurring ZIKV E proteins. Among naturally occurring E proteins, only a single amino acid is observed at most sites (Fig 3A), with a small minority of sites having two different amino acids observed in naturally occurring sequences (e.g., site 120 in Fig 3A). As a result, the correlation between our experimentally measured amino-acid preferences and amino-acid frequencies in nature was modest although still highly significant (Fig 3B). In particular, it is obvious from Fig 3B that although our experiments measure a continuum of amino-acid preferences spanning from 0 to about 0.8, the frequencies of amino acids among natural E protein sequences are usually either nearly zero or nearly one at each site.

**FIG 3:**
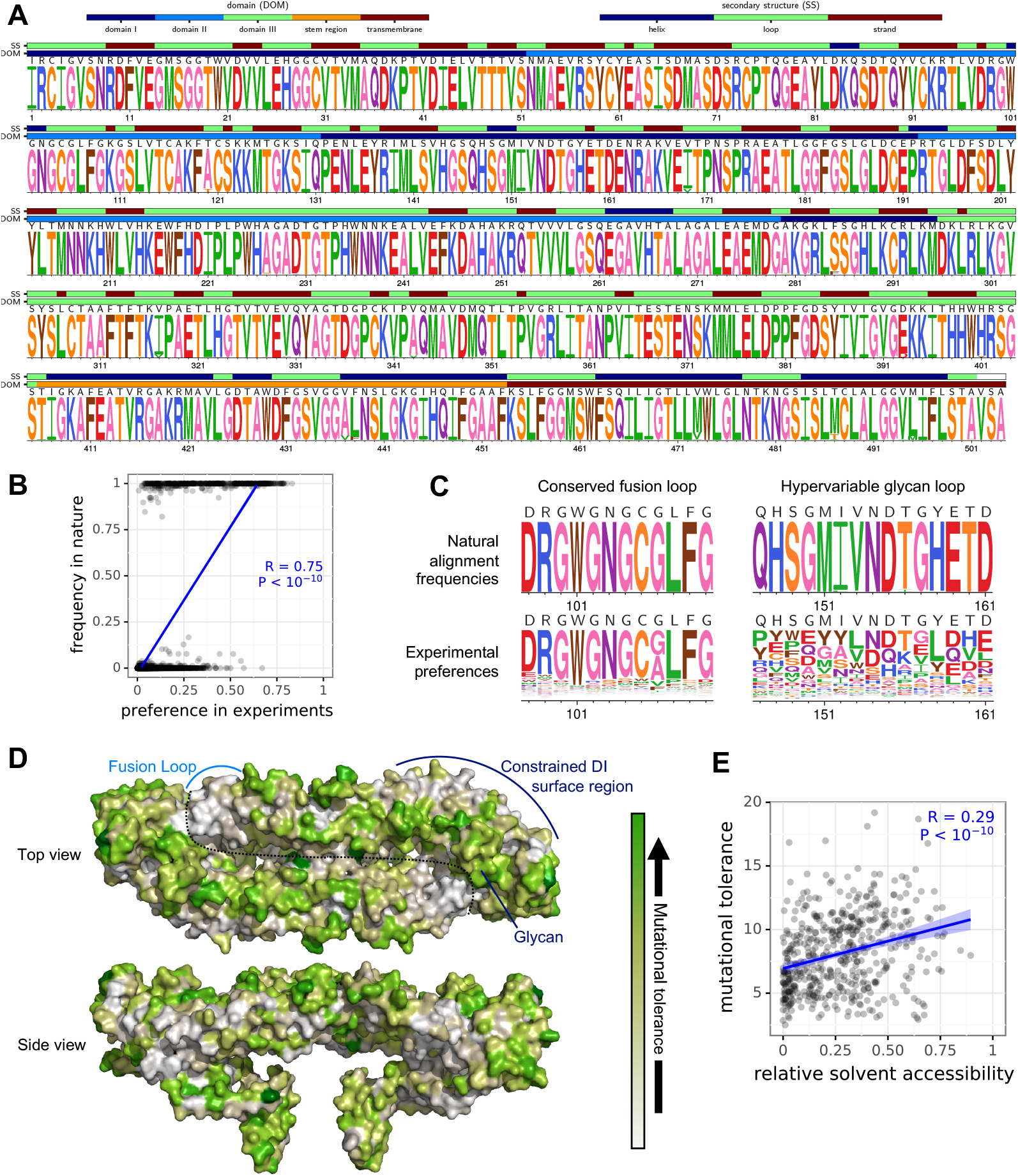
Comparison of deep mutational scanning measurements to variation in nature. **(A)** The frequencies of amino acids at each site in an alignment of naturally occurring E protein variants. The height of each letter is proportional to the frequency of that amino acid in the sequence alignment. **(B)** Correlation between amino-acid preferences measured in the experiments and the frequencies of amino acids in the natural alignment. **(C)** Direct comparison of natural aminoacid frequencies and experimentally measured amino-acid preferences in the conserved fusion loop and hypervariable glycan loop as defined in Dai et al.^26^. **(D)** Experimentally measured mutational tolerance mapped on the structure of E protein in PDB 5ire^7^. The mutational tolerance of each is quantified as the Shannon entropy of the amino-acid preferences and colored according the indicated scale. An E protein dimer is shown, with each monomer separated by a dotted line. Key features are marked, including the glycan and highly constrained fusion loop and dimer distal DI region. **(E)** Correlation between the relative solvent accessibility and mutational tolerance of each site. Relative solvent accessibility is computed using a single monomer of PDB 5ire with DSSP^48^, normalizing to the absolute solvent accessibilities in Tien et al.^49^. Mutational tolerance is computed as the number of effective amino acids, which is the exponential of the Shannon entropy of the preferences.

There are two possible reasons why the natural amino-acid frequencies are usually at the extremes of zero or one: either ZIKV E protein is intolerant of mutations in nature, or natural evolution has not sampled many of the tolerable changes. The latter explanation seems plausible because the vast majority of ZIKV sequences were collected in just the last few years from closely related strains. To distinguish these possibilities, we compared two different regions of E protein that, based on structural considerations and comparison to other flaviviruses, are thought to have wildly different levels of functional constraint: the conserved fusion loop and the hypervariable glycan loop^26^. Both regions exhibit very little variation among natural ZIKV sequences, but differ greatly in the mutational tolerance measured in our deep mutational scanning (Fig 3C). Our experiments suggest that most sites in the fusion loop strongly prefer the single amino acid that is most common in nature, but that most sites in the hypervariable glycan loop can tolerate many amino acids. These results support the notion that the high conservation of E in nature is due at least in part to the limited sampling of evolutionary space among sequenced isolates, and suggest that the deep mutational scanning is indeed a useful alternative to measure the inherent mutational tolerance of each site.

We therefore used the experimentally measured amino-acid preferences to calculate the mutational tolerance of each site in E protein, and mapped these mutational tolerance values on the protein’s structure (Fig 3D). Mutational tolerance varies widely across the protein. At the most general level, sites on the surface of E protein tend to be more mutationally tolerant than sites in the core—a trend that has also been observed for many other proteins^27–29^. To quantify this trend, we computed the correlation between a site’s solvent accessibility and mutational tolerance, and found that the correlation was modest but highly significant (Fig 3E). Strikingly, nearly the entire outward facing surface of the E protein demonstrated a high degree of mutational tolerance. Exceptions included the fusion loop region, which as pointed out above, tolerated nearly no changes and the dimer distal area of the DI region, which also appeared intolerant to mutation (Fig 3D).

### Validation of deep mutational scanning measurements with growth curves of individual viral mutants

To validate that the high-throughput deep mutational scanning measurements actually reflected mutational effects on viral growth, we generated a subset of E protein mutants and tested their growth individually. To do so, we cloned each mutation into the parental plasmid, produced homogenous stocks of these viruses by transfection of 293T cells, and then performed multicycle growth curves starting at low multiplicity of infection (MOI) in Vero cells as described in^25,30^ (Fig 4A).

**FIG 4:**
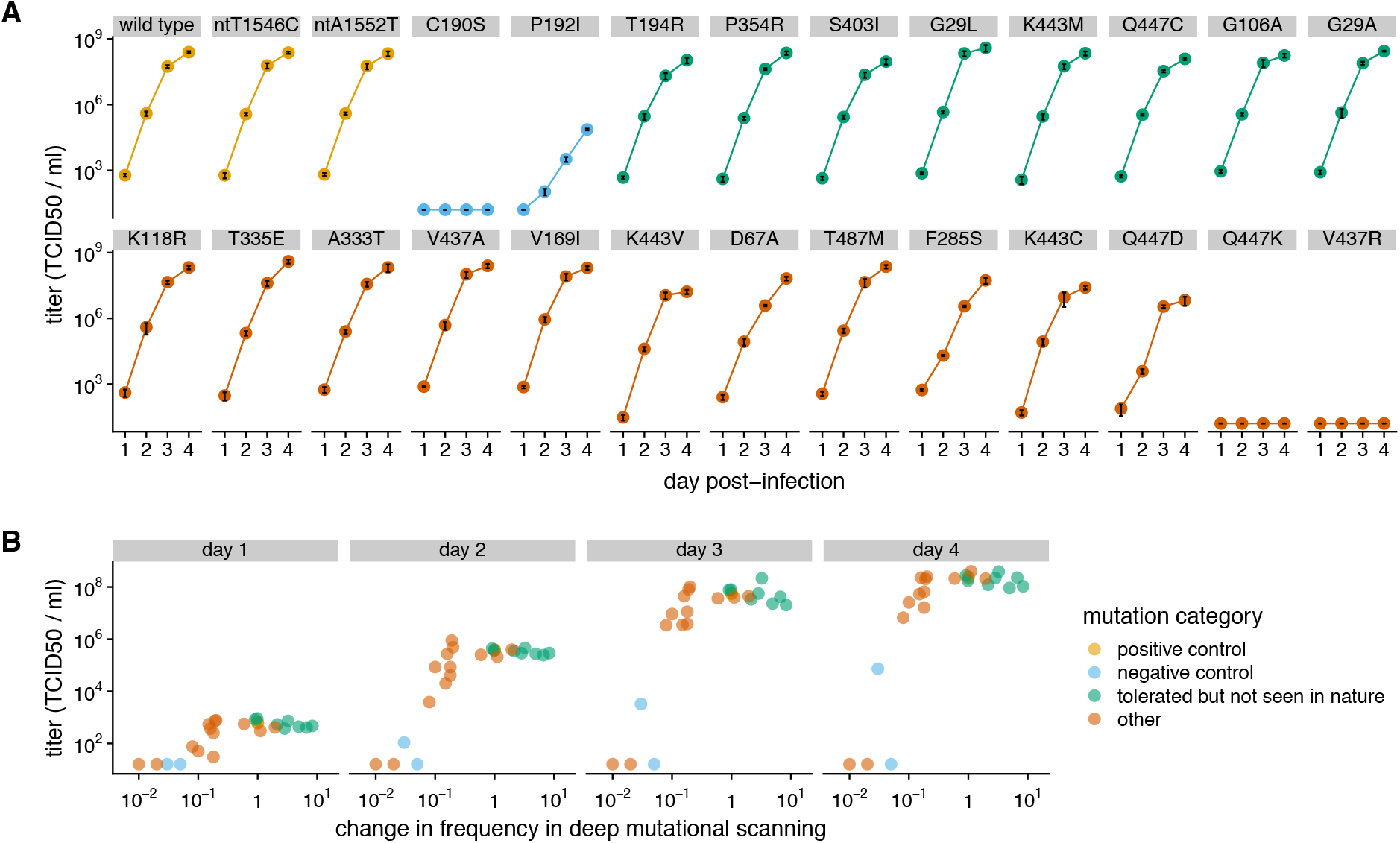
Growth curves of selected mutant viruses. **(A)** Vero cells were infected with indicated mutant virus at a MOI of 0.01, and virus in the supernatant was titered at 1, 2, 3, and 4 days post-infection. The growth curves were performed in triplicate, and the black lines inside the points show standard error. Mutant categories, as described in the results section, are indicated by marker color. **(B)** Relationship among the effect of each mutation in the deep mutational scanning and its titer in the growth curves grouped by day post infection. The x-axis represents the fold-change in frequency during the deep mutational scanning, which is the ratio of the amino-acid preference of the mutant amino acid relative to the wildtype.

For these growth curves, we selected several groups of mutants. As “positive” controls we used the wild-type virus and two synonymous mutants that are not expected to affect viral fitness — and all of these viruses grew well (Fig 4A). As negative controls, we selected a C190S mutation, as this cysteine residue is absolutely conserved in all flavivirus sequences, and P192I, as this residue is a proline in all but the most divergent tick-borne encephalitis and yellow fever viruses, where it is a valine. Indeed, C190S produced no detectable virus, and P192I was highly impaired (Fig 4A). We also selected eight mutations that our deep mutational scanning suggests should be well tolerated, but that are not observed among natural ZIKV E protein sequences. All eight of these mutants grew well (Fig 4A), supporting the notion that the deep mutational scanning can identify well-tolerated mutations that are not observed among natural sequences. We generated 13 additional mutants that were expected to (and did) have a range of effects on viral growth (Fig 4A).

Overall, these growth curves showed a good relationship between the measurements in the deep mutational scanning and the growth of the individual mutants in Vero cells (Fig 4B). Specifically, mutations that decreased by >10-fold in the deep mutational scanning strongly impaired viral growth, whereas mutations that had wild-type or better scores in the deep mutational scanning always grew well in the growth curves — even if these mutations are not observed among natural sequences. However, mutations that appeared to be just mildly attenuated in the deep mutational scanning (selected at a frequency within 10-fold of wild type) often grew about as well as wild type, suggesting that there was some noise in the high-throughput measurements and caution should be used in calling mutations as deleterious if they are only mildly depleted in the high-throughput experiments.

### Mapping mutations that escape neutralizing antibodies

Deep mutational scanning can be extended to completely map viral mutations that escape antibody neutralization in a technique termed mutational antigenic profiling^18,24^ (see Fig 5A for an outline of this approach). To demonstrate that our MR766 libraries can be used in this manner, we selected our libraries with two previously described patient-derived antibodies that efficiently inhibit ZIKV infection. One antibody, ZKA64 was shown to bind a recombinant form of domain III of the E protein, while the other, ZKA185, was originally described as a neutralizing, nonbinding antibody because it did not interact with purified full length E protein or the EDIII domain^14^.

**FIG 5:**
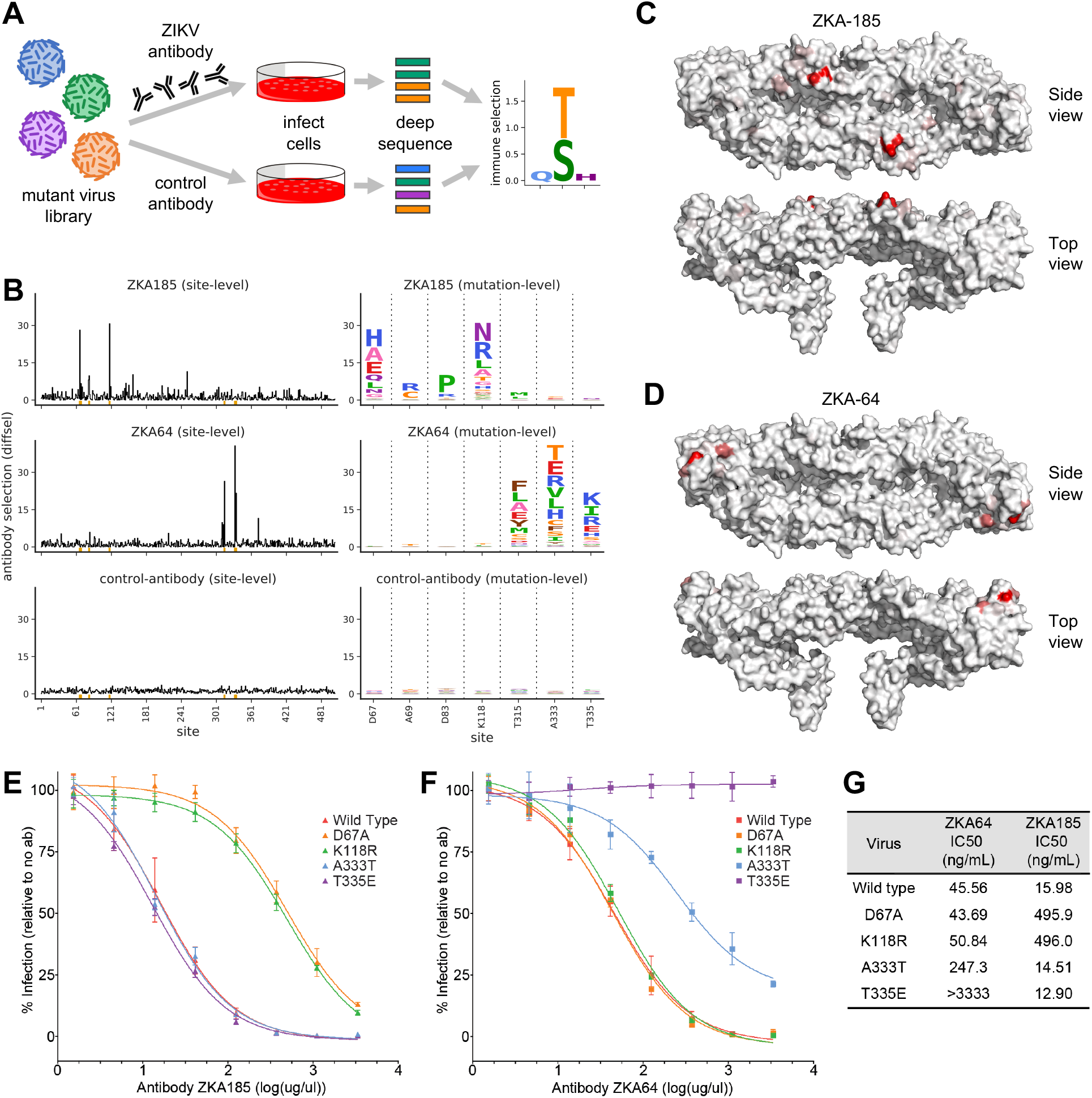
Mutational antigenic profiling to map antibody escape mutations. **(A)** Schematic of the workflow. The mutant virus libraries were incubated with ZIKV neutralizing antibodies or an irrelevant antibody, infected into cells, and the resulting viral RNA was deep sequenced to identify which mutations survive immune selection. The data can be analyzed to compute the differential selection for each mutation, which is the logarithm of its enrichment in the immune selection relative to a mock-selection control. Hypothetical results are shown, with larger letters indicating more strongly selected mutations. **(B)** The overall selection at each site (line plots) and the selection for specific mutations at strongly selected sites (logo plots) for two anti-ZIKV antibodies ZKA185 and ZKA64, as well as the negative control antibody that does not target ZIKV. **(C), (D)** The overall strength of immune selection at each site mapped on the structure of E protein, with red coloring indicating residues with differential mutant selection. **(E), (F)** Neutralization curves validated that strongly selected mutations indeed greatly reduced viral neutralization for each antibody. **(G)** IC50 values computed from the neutralization curves.

We began with the virus libraries that had been passaged at a low MOI to ensure a link between genotype and phenotype. We incubated these mutant virus populations for one hour with a concentration of antibody that neutralized roughly 99.9% of the infectious titer of wild type virus. This quantity of antibody leads to strong signal under the assumption that about 0.5 to 1% of all possible amino-acid mutations mediate escape (this is expected if the typical antibody epitope contains 2 to 8 residues, and about half the amino acid substitutions at these sites mediate escape). We then added this antibody-virus mixture to Vero cells and incubated for 24 hours. We next washed to remove unbound input virus, extracted total RNA from infected cells, and reverse-transcribed and amplified the E protein gene for deep sequencing. We simultaneously performed a control mock-neutralization with the hepatitis C virus neutralizing AR3A monoclonal antibody, which has no impact on ZIKV infection. Quantification of the viral RNA remaining after neutralization, by qRT-PCR, indicated that 1.5% to 2.6% of the mutant virus pool escaped antibody neutralization (data not shown).

Each ZIKV E antibody strongly selected mutations at just a handful of sites in E protein, whereas the control antibody did not select any mutations (Fig 5B). The different anti-ZIKV antibodies selected distinct mutations, suggesting that the mutations directly impact the antibody epitope rather than altering E protein conformation to generally increase neutralization sensitivity. We mapped the sites of strongly selected mutations onto the structure of E protein and found the most strongly selected mutations clustered near each other (Fig 5C). The ZKA64 selected mutations were all found in the DIII domain of the E protein, as previously predicted^14^. One of these residues, T315, is in the ABCD sheet epitope, and the A333 and T335 residues are in the lateral ridge epitope. Conversely, ZKA185 selected mutations all occurred at sites in the DII E protein domain. One of these sites, D67, was recently shown^31^ to be a location for a single ZKA185 resistance mutation, D67E, which our results confirm.

To validate the antibody-escape mutants, we chose two mutations expected to strongly affect sensitivity to each anti-ZIKV antibody (A333T and T335E for ZKA64, and D67A and K118R for ZKA185). We performed neutralization assays against viruses engineered to contain these mutations, using a previously described approach^14,22,32^. As expected from the mutational antigenic profiling, each mutation greatly reduced neutralization sensitivity to the antibody that selected it, but not to the other antibody (Fig 5E,F,G). These results show that mutational antigenic profiling with our viral libraries can accurately map antibody escape mutations.

## DISCUSSION

We have completely mapped the effects of all amino-acid mutations to the ZIKV E protein on viral growth in cell culture. The resulting sequence-function map can be used to assess the probable effect of every possible amino-acid mutation — something that is valuable both for understanding protein function and assessing the probable impact of mutations observed in natural sequences.

Some aspects of this sequence-function map can be interpreted in light of what is already known about E protein’s structure and function. For instance, disulfide bonds often play an important role in stabilizing proteins. Consistent with the structural importance of these bonds, our results show that the first 12 cysteines in the E protein, all of which form disulfide bonds^7^, are highly intolerant of change to any other amino acid. In contrast, the last cysteine in the protein’s sequence (C488) is not involved in a disulfide bond, and this site is highly tolerant of mutations. Similarly, ioniziation of histidine residues is important for the low pH conformational change in E protein that helps mediate membrane fusion. There are 16 histidines in the ZIKV MR766 E protein, and our map shows that five of them were strongly selected during the deep mutational scanning (H144, H214, H249, H288, and H446). These five histidines are all ones that studies of other flavivirus E proteins have suggested are important for fusion: mutating H249 and H288 led to a loss of infectivity in yellow fever virus^33^, while West Nile virus was strongly impaired by mutating H144 and H446 and more moderately impaired by mutating H214 and H446^34^. We can also rationalize some of the sequence-function map in more general terms. For instance, sites on the surface of E protein tend to be more mutationally tolerant than sites in the core, a trend that has also been observed for many other proteins^27,28^ and presumably reflects the increased constraints on residue packing in the interior of a protein. In addition, certain functionally important regions of the protein are relatively mutationally intolerant despite being on the core. These regions include the fusion loop, and a surface region extending from near the end of domain I to the beginning of domain III that has been implicated as a flavivirus receptor-binding site^12,13^.

But the greatest value of our sequence-function map is not that it confirms the expected selection at the handful of E protein residues known to be important for some specific biological function, but rather that it provides a means to assess the likely impact of the vast numbers of mutations for which there is not prior functional information. The relationship between structure and function is complex, and furthermore mutations to a protein can affect a myriad of E protein traits ranging from translation to folding to assembly to entry. Using our map, we can make observations about mutational effects that are virtually impossible to rationalize from current molecular knowledge: for instance, we can hypothesize no clear explanation of the constraints at site 447 that explain why the virus is completely tolerant of Q447C, is substantially attenuated by Q447D, and is completely inactivated by Q447K — yet both our map and our validation experiments show that this is the case. Because the effects of many other mutations are similarly difficult to rationalize, our map forms a useful reference for studies of E protein evolution and function.

The key to the technical success of our experiments was the use of a highly efficient plasmid-based reverse-genetics system to generate the mutant virus library^25^. Flavivirus genomes are notoriously difficult to handle in traditional plasmid-based cloning systems since they often contain nucleotide sequences that are toxic in bacteria. The plasmid used in our experiment has an intron inserted into the most toxic region, which eliminates the bacterial toxicity but spliced in mammalian cells and so does not affect the actual generated virus^25^. By using this plasmid and also choosing a viral strain (MR766) that grows to high titer in cell culture, we were able to generate viral libraries that effectively sampled all amino-acid mutations to E protein. This library diversity contrasts to two other recent ZIKV deep mutational scanning studies^21,22^: both of these studies revealed important biological insights about host adaptation, but neither produced a complete map of amino-acid level selection due to an inability to generate and maintain the viral library diversity to sample all amino acids. Of course, our experiments were still subject to noise, probably because even our high-efficiency reverse-genetics had some bottlenecking of viral diversity. Our validation experiments with single-mutant growth curves suggested that mutations that are strongly depleted in the deep mutational scanning (decrease >10-fold, equivalent to a mutational effect < log2 0.1 = −3.3 in raw data provided in Supplementary File 3) are consistently deleterious for viral growth. Likewise, mutations that the deep mutational scanning suggest are comparable preferred to wildtype are consistently well tolerated. However, caution should be used in interpreting the effects of mutations that the deep mutational scanning suggests are just mildly deleterious, since these modest effects fall on the boundary between the signal and noise of the high-throughput measurements.

A major point to keep in mind is that our study mapped how mutations affect the growth of a single strain of ZIKV in Vero cells, meaning that there are two major caveats that must be kept in mind when interpreting the results. First, the MR766 ZIKV strain we used was isolated in 1947 and was subsequently passaged in the lab^10^. Therefore, the effects of some mutations could differ between this strain and more contemporary ZIKV strains. Indeed, prior deep mutational scanning studies of other viruses have shown that mutational effects do shift somewhat among different strains of the same virus^19,35,36^, although these shifts tend to be relatively small when the strains are closely related as is the case for all known ZIKV strains. Second, Vero cells are a relatively permissive monkey-derived cell line, whereas actual ZIKV evolves under selection to efficiently replicate in actual humans and mosquitoes. Indeed, prior deep mutational scanning studies of ZIKV^21,22^ and other viruses^37,38^ have identified numerous mutations with effects that depend on the host species and level of innate-immune function in the cells used to grow the virus. Therefore, our map should be interpreted as providing a baseline estimate of the functional effects of mutations to the E protein in a relatively permissive context.

However, the fact that the cell-culture model used in our experiment is somewhat artificial represents an opportunity as well as a caveat. In particular, it provides the capacity to directly manipulate the selective pressure to map how mutations affect some particular relevant process. In our case, we have used this fine-grained level of control to perform the first complete mapping of ZIKV antibody-escape mutants by selecting the virus library in the presence and absence of monoclonal antibodies. This work is of special interest because the conformational flexibility of E protein means that antibody neutralization mechanisms can be complex^39^, with some neutralizing antibodies not even binding to recombinant E protein^14^.

We mapped all viral mutations that affected neutralization by two antibodies, ZKA64 and ZKA185^14^. Both antibodies strongly selected mutations at a small handful of spatially proximal sites, thereby defining the functional epitopes. For ZKA64 this functional epitope was in domain III of E protein, consistent with the original report that it bound to this domain^14^. The ZKA185 antibody does not bind to recombinant E protein, so the original paper describing this antibody was unable to map its epitope^14^ although later work found a single neutralization-resistant variant with mutations at sites 67 and 84 in domain II of E protein^31^. Consistent with this later study, we found that ZKA185 exerted strong selection at a handful of sites in domain II, including site 67 and to a much lesser degree site 84. Notably, the sites selected by ZKA185 are well separated on linear E sequence, but tightly grouped on the protein structure. Overall, we think the ability to use our libraries to completely map ZIKV antibody-escape mutations could prove very useful. Antibody-escape mutants are traditionally isolated by passaging virus in the presence of neutralizing antibody, but each such experiment typically only isolates one or a few of the possible escape mutants. In contrast, our experiments completely map all amino-acid mutations that affect neutralization, and so provide a full picture of the functional epitope.

Overall, our work provides a complete map of how E protein amino-acid mutations affect viral growth in a permissive cell line, and viral escape from two monoclonal antibodies. This map in and of itself is a valuable complement to structural maps of E protein, and will be useful for interpreting and designing mutagenesis studies to probe the function and antigenicity of the protein. In addition, our work provides a template for future studies to map mutations that affect more specific viral phenotypes, such the ability to replicate in specific cell lines or animals / mosquitos, or escape from antigenic recognition by other antibodies or even polyclonal sera.

## MATERIALS AND METHODS

### Cell lines and culture

293T (provided by Charles M. Rice, Rockefeller University) and Vero (ATCC CCL-81, Manassas, VA) cells were grown in Dulbecco’s modified Eagle’s medium (DMEM; Gibco BRL Life Technologies, Gaithersburg, MD) with 10% fetal bovine serum (FBS; Gibco BRL Life Technologies).

### Antibodies

ZKA185 and ZKA64 are human monoclonal neutralizing antibodies targeting ZIKV E protein^14^ (Ab00835-15.0 and Ab00779-15.0, respectively, Absolute Antibody, Oxford, UK). The humanized monoclonal antibody D1-4G2-4-15 (4G2) (Absolute Antibody, Oxford, UK) is a broadly reactive flavivirus antibody that binds to an epitope at the fusion loop domain of the E protein^40^. Rabbit anti-NS3 was raised against a peptide consisting of amino acid residues 456 to 469 of the ZIKV MR766 sequence (GenBank accession number <u>AAV34151</u>). The anti-HCV E2 monoclonal antibody (clone AR3A) used as control for the mutational antigenic profiling was provided by Mansun Law (Scripps Research Institute)^41^.

### Creation of mutant plasmid libraries

The mutant plasmid libraries were cloned into our previously described single-plasmid reverse genetics system for ZIKV strain MR766^25^ (sequence is available at Genbank accession KX830961). In order to enable efficient cloning into this plasmid backbone, we designed a recipient plasmid in which a 1.65 kb region of this plasmid including the entire E protein coding sequence was replaced by eGFP flanked by NotI sites, and named this plasmid ZIKV_MR766_int_GFP (the “int” stands for the fact that the clone contains an intron to reduce toxicity in bacteria^25^). The sequence of ZIKV_MR766_int_GFP in Genbank format is at https://github.com/jbloomlab/ZIKV_DMS_with_EvansLab/blob/master/data/1726_ZIKV_MR766_int_GFP.gb.

We created codon-mutant libraries of the E gene using a previously described PCR mutagenesis approach^23^. We first designed mutagenic primers that tiled all codons in the E gene. These primers were designed using the computer scripts at https://github.com/jbloomlab/CodonTilingPrimers to have an average melting temperature of about ~65°C as in^24^. The primer sequences are in Supplementary File 1. We performed the mutagenesis PCR using these primers and the following end primers flanking E gene: 5’-gccattgcctggcttttgggaagc-3’ and 5’-tggtacttgtaccggtccctccaggc-3’. Two rounds of mutagenic PCR were performed in triplicate exactly as in^23^. After NotI digestion and purification of the recipient plasmid backbone ZIKV_MR766_int_GFP, mutagenized E gene amplicons were cloned into the recipient MR766 genome using Gibson assembly and transformed into 10-beta electrocompetent *E. coli* (NEB). Transformants were plated on LB supplemented with carbenicillin. Each of the triplicate libraries contained >1×10^6^ unique transformants as estimated by plating a dilution of the library transformations. Fifteen individual colonies of each library were Sanger sequenced and summarized in Fig.1. Each codon mutant library was prepared by HiSpeed Maxiprep (Qiagen) a pool of plated transformants resuspended in LB. In order to help ensure higher plasmid stability, the bacterial growth steps were performed at 30°C.

### Cloning single mutant ZIKV

Individual amino acid mutants were generated by PCR, using previously described techniques^25,42^. Briefly, two round of PCR were performed. The first round involved two reactions: one reaction with a forward oligo (sequences available upon request) that overlapped unique restriction site in the ZIKV strain MR766 plasmid^25^ and a reverse primer that encodes the desired mutation, and the second reaction with a forward oligo also bearing the desired mutation and a universal reverse oligo the spanned another unique restriction site. These products were then combined as template in a third PCR reaction with only the outside universal oligos. This product was cloned in the parental plasmid using the In-Fusion Cloning Kit (Clontech, Mountain View, CA). All PCR amplified sequences, cloning sites, and desired mutations were verified by Sanger sequencing.

### Rescue of ZIKV E libraries, WT or single mutant ZIKV

Viruses and ZIKV E libraries were produced by transfecting plasmids into 293T cells as previously reported^25,32^. Briefly, prior to transfection, 6-well plates and 24-well plates were seeded overnight with 1.2 × 10^6^ or 0.3 x 10^6^ cells/well respectively. For each library replicate, 6 wells of a 6-well plate were transfected using 1 μg of plasmid DNA per well and the *Trans*IT-LT1 transfection reagent (Mirus Bio, Madison WI) per the manufacturer’s recommendations. The viral supernatants were collected at day 2 posttransfection, pooled, filtered (pore size, 0.45 μm) to remove cellular debris, and stored at −80°C. For each defined mutant virus, 1 well of a 24-well plate was transfected using 1 μg of plasmid DNA per well and the *Trans*IT-LT1 transfection reagent (Mirus Bio, Madison WI) per the manufacturer’s recommendations. The viral supernatants were collected at day 2 posttransfection, filtered (pore size, 0.45 μm) to remove cellular debris, and stored at −80°C.

### Titration of infectious viruses and ZIKV E libraries

For screening, the infectious titers of rescued wild type, as a positive control, and mutant libraries were determined by cytopathic effect (CPE)-based limiting dilution assay on Vero cells, based on techniques previously described^25,32^. One day prior to infection, 96-well plates were seeded at a density of 10^4^ cells/well. They were infected with 100 μl of virus serially diluted in DMEM with 2% FBS (8 wells per dilution). Infection was scored by detection of ZIKV-induced CPE, and TCID50 units were calculated according to the method of Reed and Muench^43^. Individually rescued mutants were titered by immunostaining with the pan-flavivirus E protein reactive 4G2 antibody and flow cytometry, as previously described^32^. Briefly, one day prior to infection, 24-well plates were seeded at a density of 1.25×10^5^ cells/well. The next day, 250 uL of serially diluted transfection supernatants in DMEM with 2% FBS was added to each well (in triplicate). Staining was performed at 24 hours post infection to avoid any spread. Only the viral dilutions leading to 30 to 1% of infected cells were used to calculate the infectious titers. Infectious unit per mL were calculated using the following formula: “percentage of infectious events x number of cells / volume (mL) of viral inoculum”.

### Selection of ZIKV E libraries and control WT ZIKV

Ten 15 cm-diameter petri dishes, per replicate or WT control, with 10^7^ cells on the day of infection, were infected at a MOI of 0.01 infectious units per cell. The viral supernatant was collected at day 2 and 3 post infection, filtered (pore size, 0.45 μm) and stored at −80°C. Infected cells were collected at day 3 post infection, total RNA were extracted using the RNeasy Plus kit (Qiagen, Valencia, CA) as per manufacturer recommendation.

### Growth curves

Vero cells (1 × 10^5^) were seeded in a well of a 24-wells plate, 1 day prior to infection, in triplicate, and then infected at a MOI of 0.01 infectious units per cell in DMEM with 2% FBS. Virus was removed 6 hours later, and cells were washed twice with PBS before 1 mL of DMEM with 2% FBS was added. Supernatants were collected and filtered (0.45 μm pore size) daily for the next 4 days. Supernatant infectivity was determined by TCID50 assay as described above.

### Measurement of the inhibitory concentration 99.9 (IC99.9)

24-well plates were seeded one day prior to infection at a density of 1.25 x 10^5^ cells/well in 500 μl DMEM supplemented with 2% FBS. 2-fold serial dilutions of antibody were incubated, at 37°C for one hour, with 2.5 x 10^5^ infectious unit of control WT virus “selected” on Vero cells, in triplicate in 500 uL media. Culture media on the above Vero cells was then replaced with the virus-antibody dilutions for 24 hours. Cells were collected after extensive washes with PBS. Total RNA was extracted with the RNeasy Plus kit (Qiagen). Viral RNA quantification was performed on 20 ng of RNA by ZIKV-specific quantitative RT-PCR, as described below. IC99.9 was calculated using Prism software (GraphPad Software) (dose-response, three parameters) for each antibody. For ZKA64 and ZKA185 the IC99.9 values were 531 and 274 ng/mL, respectively.

### Selection of antibody-escape mutants

One day prior to infection, 6-well plates were seeded at a density of 5 x 10^5^ cells/well in 2 mL DMEM supplemented with 2% FBS. Both ZKA64 and ZKA185, at the IC99.9 concentration stated above, and AR3A at 1 ug/mL, were incubated, at 37°C for one hour, with 10^6^ infectious units of ZIKV E selected libraries or control WT virus in 2 mL of DMEM with 2% FBS. Each antibody-mixture was then added to the cells. At 24 hours post-infection, cells were extensively washed to remove the inoculum and RNA was extracted from the cells using RNeasy Plus kit (Qiagen) for deep sequencing.

### Neutralization assays

One day prior to infection, 24-well plates were seeded with 1.25×10^5^ Vero cells per well. Threefold serial dilutions of antibodies were incubated with 2.5×10^4^ (MOI, 0.1) infectious units of virus at 37°C for 30 min and then used to infect Vero cells. At 24 h post-infection, cells were collected and analyzed by flow cytometry, as described above, to determine the percentage of infected cells. The percentage of infected cells in each well was normalized relative to the percentage of infected cells in the absence of antibody. Analysis of dose-response curves was performed with Prism software (GraphPad Software) (dose-response, three parameters) to calculate the apparent IC50 for each antibody.

### ZIKV specific quantitative RT-PCR

Viral RNA quantification was performed by ZIKV-specific quantitative RT-PCR, as previously described^25^. Total RNA was extracted with the RNeasy Plus kit (Qiagen, Valencia, CA) as per manufacturer’s recommendation. For quantifying ZIKV RNA, 20 ng of total RNA was *in vitro* reverse-transcribed using the High-Capacity cDNA Reverse Transcription Kit (Thermo Fisher Scientific, Waltham MA). Quantitative PCR was performed with ZIKV-specific primers (5′ TTGGTCATGATACTGCTGATTGC and 5′ CCYTCCACRAAGTCYCTATTGC) and the LightCycler 480 SYBR Green I Master mix (Roche Applied Science) using the LightCycler 480 II real-time PCR system (Roche Applied Science). Sequences of the primers targeting ZIKV were adapted from previously published sequences^44^. Quantification of ZIKV RNA copies per ug of total RNA was performed against a standard curve of *in vitro*-transcribed MR766 ZIKV RNA using the T7 RiboMAX™ Express Large Scale RNA Production System (Promega).

### Flow cytometry

Cells were fixed in 4% paraformaldehyde and stained with 4G2 (E protein) or NS3 antibodies and goat-anti-human or donkey anti-rabbit secondary antibodies, respectively, conjugated to Alexa Fluor 647 (Thermo Fisher Scientific, Waltham MA) using methods previously described^45^. Data were acquired on a Fortessa flow cytometer (Becton, Dickinson, Franklin Lakes, NJ) and analyzed using FlowJo software (Tree Star, USA).

### Deep sequencing using barcoded-subamplicon sequencing

For each sample, viral RNA was extracted with the RNeasy Plus kit (Qiagen, Valencia, CA) as per manufacturer’s recommendation. The E gene was then reverse transcribed using AccuScript Reverse Transcriptase, and primers flanking the E coding sequence (5’-gccattgcctggcttttgggaagc-3’ and 5’-tggtacttgtaccggtccctccaggc-3’).

To obtain high sequencing accuracy, we used the barcoded-subamplicon sequencing described in^17^ (see also https://jbloomlab.github.io/dms_tools2/bcsubamp.html). We generated five subamplicons of approximately 430 bp that tile the length of the E coding sequence. Each primer used for amplification adds an N8 randomized barcode and Illumina adapter sequences. These primers are provided in Supplementary File 2. Round1 PCR reactions were set up using 12 μL of 2× KOD Hot Start Master Mix, 4 ng of cDNA template, 2 μL of forward primer diluted to 5 μM, and 2 μL of reverse primer diluted to 5 μM. The cycling program used for the Round1 PCR was: 1) 95°C for 2:00; 2) 95°C for 0:20; 3) 70°C for 0:01; 4) 50°C for 0:20; 5) 70°C for 0:20; 6) Go to (2) (9 times); 7) 95°C for 1:00; 8) 4°C hold. The final denaturation step ensures that double-stranded DNA molecules entering Round2 PCR will contain two uniquely barcoded variants.

Subamplicons were purified using AMPure XP beads (1:1 bead-to-sample volume), diluted, and used as template for a second round of PCR using ~8×10^5^ single-stranded template molecules per subamplicon, per sample. The primers used for Round2 PCR add sample-specific indices and the Illumina cluster-generating sequences. The Round2 PCR reactions contained 20 μL of 2× KOD Hot Start Master Mix, Template DNA diluted to 8×10^5^ ssDNA molecules per subamplicon, 4 μL of forward primer diluted to 5 μM, and 4 μL of reverse primer diluted to 5 μM. The cycling program used for Round2 PCR was: 1) 95 °C for 2:00; 2) 95°C for 0:20; 3) 70°C for 0:01; 4) 55°C for 0:20; 5) 70°C for 0:20; 6) Go to (2) (29 times); 7) 4°C Hold. The Round2 subamplicons were pooled, size selected by gel extraction (Zymo) and subsequently purified with AMPure XP beads. Subamplicons were sequenced with an Illumina HiSeq2500 using 2 × 250 bp paired-end reads in Rapid Run mode.

The raw deep sequencing data have been deposited in the Sequence Read Archive as BioProject PRJNA530795 (https://www.ncbi.nlm.nih.gov/bioproject/PRJNA530795).

### Analysis of deep sequencing and data availability

Computer code that performs all of the analyses of the deep sequencing data is on GitHub at https://github.com/jbloomlab/ZIKV_DMS_with_EvansLab. This repository includes a notebook that gives detailed statistics on read depth, mutation frequencies, etc for all samples (see https://github.com/jbloomlab/ZIKV_DMS_with_EvansLab/blob/master/results/summary/analysis_notebook.md).

Briefly, we used dms_tools2^46^ (https://jbloomlab.github.io/dms_tools2/), version 2.4.14, to process the deep sequencing data to count the occurrences of each mutation in each sample (see https://jbloomlab.github.io/dms_tools2/bcsubamp.html for details on how the software does this). The amino-acid preferences were computed from these counts using the approach described in^46^ (see also https://jbloomlab.github.io/dms_tools2/prefs.html). The mutational effects are simply the log of the preference for the mutant amino acid divided by the preference for the wild-type amino acid. The differential selection in the antibody selections was computed using the approach described in^18^ (see also https://jbloomlab.github.io/dms_tools2/diffsel.html), computing selection for each antibody relative the no-antibody control condition. The figures in the paper show just the positive differential selection. For both the amino-acid preferences and antibody differential selection, the paper reports the average value of the three experimental replicates. To make the logo plots of the antibody selection in Fig 5B, we used the program dmslogo, version 0.2.3^47^.

The raw numerical values of the counts of mutations in each sample, the amino-acid preferences, and the differential selection are provided in CSV files at https://github.com/jbloomlab/ZIKV_DMS_with_EvansLab. In addition, Supplementary File 3 gives these numerical values in Excel format.

### Sanger sequencing of passaged viruses

We verified that the sequence of individually rescued mutant viruses that displayed impaired growth characteristics were stable throughout the course of the above experiments. To do so, total RNA from 100 uL of supernatant from the day 4 post-infection growth curve was extracted with the EZNA Viral RNA Kit (Omega Bio-Tek), as per manufacturer’s recommendation. The extracted RNA was used as a template for random hexamer-primed cDNA synthesis using the SuperScript III First-Strand synthesis system (Thermo Fisher Scientific, Waltham, MA). This cDNA was used for PCR using the Expand High-Fidelity PCR system (Roche Life Sciences, Indianapolis, IN) with oligonucleotides flanking each mutant region and bulk PCR products subjected to Sanger sequencing. No reversions or otherwise unintended mutations were found in any virus following the growth curve experiments.

### Structural analyses

Mutational tolerance and antibody escape were mapped onto the surface representation of the E protein dimer of PDB structure 5IRE using PyMol. Mutational tolerance was quantified by the Shannon entropy on a scale from white (lower entropy, less mutational tolerance) to green (higher entropy, higher mutational tolerance). Sites of escape from monoclonal antibodies were mapped on a gradient according to the positive site differential selection (averaged across triplicates) and colored on a spectrum from white (low selection) to red (high selection).

## Acknowledgements

We thank Adam Dingens for assistance with data analysis, Charles Rice (Rockefeller University) for 293T cells, and Mansun Law (Scripps) for the HCV AR3A negative control antibody. We thank the Fred Hutch Genomics Core for performing the deep sequencing and the Dean’s Flow Cytometry CORE at the Icahn School of Medicine at Mount Sinai for FACS assistance. This work was supported by grants R21 AI133649 (to MJE and JDB), R21 AI140196 (to MJE) and R01 AI141707 (to JDB). JDB is an Investigator of the Howard Hughes Medical Institute. MJE and JDB hold Investigators in Pathogenesis of Infectious Disease Awards from the Burroughs Wellcome Fund.

---

**SUPPLEMENTARY FILE 1:** This Excel document gives the sequences of the mutagenic primers for the codon mutagenesis of the E protein. Each sheet in the document is a 96-well plate of forward or reverse mutagenesis primers.

**SUPPLEMENTARY FILE 2:** This Excel document gives the sequences of the primers used for the barcoded-subamplicon sequencing.

**SUPPLEMENTARY FILE 3:** This Excel document gives the amino-acid preferences under selection for viral growth, the mutational effects under selection for viral growth, and the mutation- and site-level selection from each antibody.

